# A physiological role for GABA_A_ receptor desensitization – induction of long-term potentiation at inhibitory synapses

**DOI:** 10.1101/2020.06.02.128900

**Authors:** Martin Field, Philip Thomas, Trevor G Smart

## Abstract

GABA_A_ receptors (GABA_A_Rs) are pentameric ligand-gated ion channels distributed throughout the brain where they mediate synaptic and tonic inhibition. Following activation, these receptors undergo desensitization which involves entry into long-lived agonist-bound closed states. Although the kinetic effects of this state are recognised and its structural basis has been uncovered, the physiological impact of desensitization on inhibitory neurotransmission remains unknown. Here we describe an enduring new form of long-term potentiation at inhibitory synapses that elevates synaptic current amplitude for 24 hrs following desensitization of GABA_A_Rs in response to prolonged agonist exposure or allosteric modulation. Using receptor mutants and allosteric modulators we demonstrate that desensitization of GABA_A_Rs facilitates their phosphorylation by PKC, which increases the number of receptors at inhibitory synapses. These observations provide a new physiological relevance to the desensitized state of GABA_A_Rs, acting as a signal to regulate the efficacy of inhibitory synapses during prolonged periods of inhibitory neurotransmission.

## Introduction

The mammalian central nervous system forms the central repository for learning and memory and is well known to exhibit structural and functional plasticity. Plasticity is usually manifest by short-or long-term alterations to the efficacy of synaptic transmission^1,2^. Historically, the most described and studied forms of neural plasticity are associated with excitatory neurotransmission and synapses^3,4^. However, over time it has become clear that these are not the only structures within the brain that exhibit structural and functional plasticity. In particular, inhibitory synapses, which are predominantly populated with GABA_A_ receptors (GABA_A_Rs) and respond to GABA released by interneurons, are increasingly recognised as highly plastic structures^5,6^. However, the mechanisms that regulate the plasticity of inhibitory synapses are incompletely understood. One important question relates to how the postsynaptic cell detects and determines the level of inhibitory synaptic activity in order to initiate appropriate forms of plasticity. Excitatory synapses possess Ca^2+^-permeable NMDA receptors which enable glutamate in the synaptic cleft to directly influence intracellular signalling in the postsynaptic cell. This results in potentiation of the synaptic response largely due to changes to the number of receptors in the postsynaptic membrane^7,8^. However, for inhibitory synapses, whether such a direct signalling link exists between synaptic activity and plasticity is unclear. The widespread occurrence of shunting inhibition in the adult CNS^9^ means that inhibitory synapse activation will often result in negligible changes to either membrane potential or transmembrane Cl^-^ fluxes.

Here we provide evidence for inhibitory synaptic plasticity being regulated by agonist-induced conformational changes to synaptic GABA_A_Rs. The most likely candidate for mediating such a link between GABA_A_R structure and the plasticity of inhibitory synapses, and thus registering their level of activity, is the desensitized state. This is a long-lived agonist-bound closed state that the receptor can enter after binding agonist and opening of the integral ion channel^10^. Recent structural work has shown that the key differences between this state and the agonist-bound open (conducting) state are conformational changes that occur at the ‘internal face’ of the receptor. This involves rotation of the receptor’s transmembrane domain α-helices and a collapse of the internal portal of the ion channel to limit the transmembrane flux of ions^11-15^.

Entry into this desensitized state occurs during the presence of bound agonist at the orthosteric site, but the process is fully reversible, and the state has a lifetime between hundreds of milliseconds to seconds^10,16^. The desensitization of ligand-gated ion channels is therefore distinct from that of metabotropic receptors in being a relatively transient process that need not involve post-translational modification of the channel or any subsequent receptor internalisation. However, this does not exclude the possibility that entry into the desensitized state can initiate long-term plasticity at inhibitory synapses.

For GABA_A_ receptors, which are members of the pentameric ligand-gated ion channel family (pLGIC) and are the major inhibitory neurotransmitter receptors in the central nervous system^9^, the long-term effects of their entry into desensitized states at inhibitory synapses have not been described. Nevertheless, many studies have focused on changes to the receptor following sustained activation with agonists or allosteric modulators^17^. These changes include: phosphorylation of receptor subunits^18-21^, altered expression of GABA_A_Rs^18,21-28^, changes to receptor mobility and clustering^26,29-31^, and alterations to receptor pharmacology^32^. In most cases, it is unclear which event triggers these long-term receptor alterations. For example, they may be a consequence of direct responses to Cl^-^ currents flowing through open GABA channels, or they may occur as a result of the cellular inhibition *per se*. However, these same conditions of prolonged or elevated GABA_A_R activation will also increase the residency of receptors in the desensitized state, which may act as a signal of GABA_A_R activity to regulate synaptic efficacy or function, as has previously been speculated^21,33^. Indeed, for the nicotinic acetylcholine receptor, prolonged application of the agonist nicotine that causes desensitization^34^, also promotes upregulation of these receptors via multiple mechanisms^35,36^. In our study using hippocampal neurons, we reveal a clear agonist-induced plasticity of synaptic GABA_A_Rs, expressed in the form of a long-term potentiation at inhibitory synapses. The reliance of this plasticity on GABA_A_R desensitization is interrogated using mutant receptor subunits that substantially alter the ability of the receptor to access the desensitized state. We further demonstrate that inhibitory long-term potentiation (iLTP) depends on the ability of receptor desensitization to promote the phosphorylation of residues that regulate the clustering of GABA_A_Rs. Overall, our data reveal a signalling mechanism whereby prolonged activation of GABA_A_Rs can lead to the strengthening of inhibition via GABAergic synapses.

## Results

### GABA induces long-term potentiation at inhibitory synapses

To assess the nature of agonist-dependent plasticity at inhibitory synapses, hippocampal neurons were exposed initially to 1 mM GABA for 20 mins. Neurons were subject to whole-cell recording after approximately 24 hrs following wash-off of GABA with control culture medium. This pre-treatment with GABA was undertaken in culture maintenance media (see Methods), rather than ACSF or similar saline solution, in order to avoid the activation of autophagic signalling pathways by nitrogen starvation, which could affect the function of inhibitory synapses^37,38^. Furthermore, cells were not subjected to electrophysiological recording during pre-treatment in order to avoid GABA_A_R rundown that is known to occur under such conditions^39-41^, and which may obscure the induction of synaptic plasticity.

GABA pre-treatment caused large (∼70%) increases in the sIPSC amplitudes, from −55.5 ± 2.0 pA in non-treated controls, to −98.7 ± 7.9 pA (Fig. 1a.b; t test: t_(7)_ = 4.75, p = 0.0021). No change was apparent in the inter-event intervals (IEIs) of the IPSCs, suggesting no concomitant alteration occurs in synaptic release probability or indeed in network activity (Fig. 1c; t test: t_(7)_ = −0.21, p = 0.83). The increase in the magnitude of the sIPSCs was evident in the distributions of both the peak sIPSC amplitudes, and the rate of rise of the synaptic events (Fig. 1d,e). Agonist-dependent plasticity is therefore evident in these neurons, driven by GABA activation, and it is expressed as an increase in postsynaptic event amplitude at inhibitory synapses.

**Figure 1.**
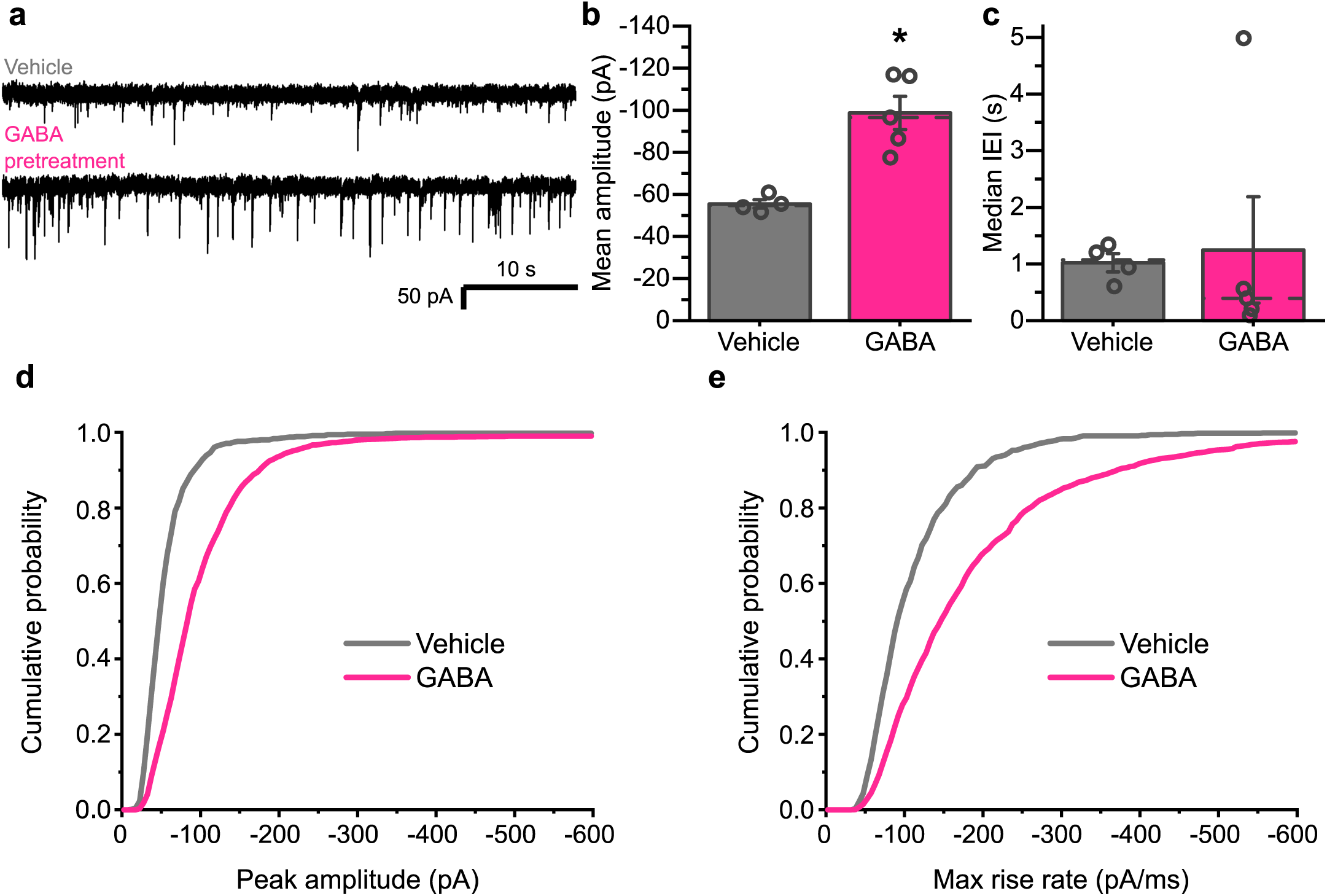
GABA causes a long-term increase in inhibitory synapse efficacy. **a** Representative spontaneous inhibitory postsynaptic currents (sIPSCs) recorded from cultured hippocampal neurons voltage-clamp at −60 mV and pre-treated with either vehicle or 1 mM GABA for 20 mins approximately 24 hrs prior to recording. **b** Bar graph of mean amplitudes of sIPSCs displaying clean, monophasic rise phases, and **c**, median inter-event intervals (IEIs) following pre-exposure to vehicle or 1 mM GABA (± S.E.M). **d** Mean cumulative probability distribution of sIPSC amplitudes and **e**, maximum rate of rise slopes for all sIPSCs recorded after vehicle or GABA treatment. n = 718 events from 4 neurons, and 2519 events from 5 neurons, respectively.

### Enhanced desensitization of GABA_A_Rs promotes inhibitory synaptic strength

To explore the role of desensitization for inhibitory synaptic plasticity, we selected previously determined mutations to synaptic GABA_A_R subunits that alter the entry of the receptor into desensitized states. These included α2^V297L^ to reduce, and γ2L^V262F^ to increase, GABA_A_R desensitization^11^. We first assessed the ability of these two mutations to alter the occupancy of the desensitized state on time scales relevant to synaptic signalling by rapidly-applying GABA to outside-out patches excised from HEK cells expressing α2β2γ2L receptors using a rapid drug application system. When exposed to 10 mM (saturating) GABA for 200 ms, the GABA currents decayed in accordance with a developing macroscopic desensitization during GABA exposure (Fig. 2a). We measured the extent of desensitisation from the depression of the steady-state GABA current relative to its peak current. The inclusion of α2^V297L^ substantially decreased the extent of this decay, from 55.6 ± 5.9 % in the wild-type (wt) to 11.7 ± 4.3 % (Fig. 2a,b; one-way ANOVA: F_(2, 17)_ = 92.45, p = 7.3×10^−10^; Tukey test (wt, α2^V297L^): p = 3.4×10^−6^), suggesting minimal accumulation of the receptor in desensitized states over this timescale for the α2 subunit mutant. Conversely, γ2L^V262F^ markedly increased the extent of GABA current decay to 82.7 ± 2.4 % compared to wild-type (Fig. 2a,b; Tukey test (wt, γ2L^V262F^): p = 0.0004), in agreement with its predicted behaviour of promoting the accumulation of GABA_A_Rs in desensitized states. Interestingly, neither mutation significantly altered the rate at which the receptor currents decayed (Fig. 2c; one-way ANOVA: F^(2,15)^ = 0.18, p = 0.84).

**Figure 2.**
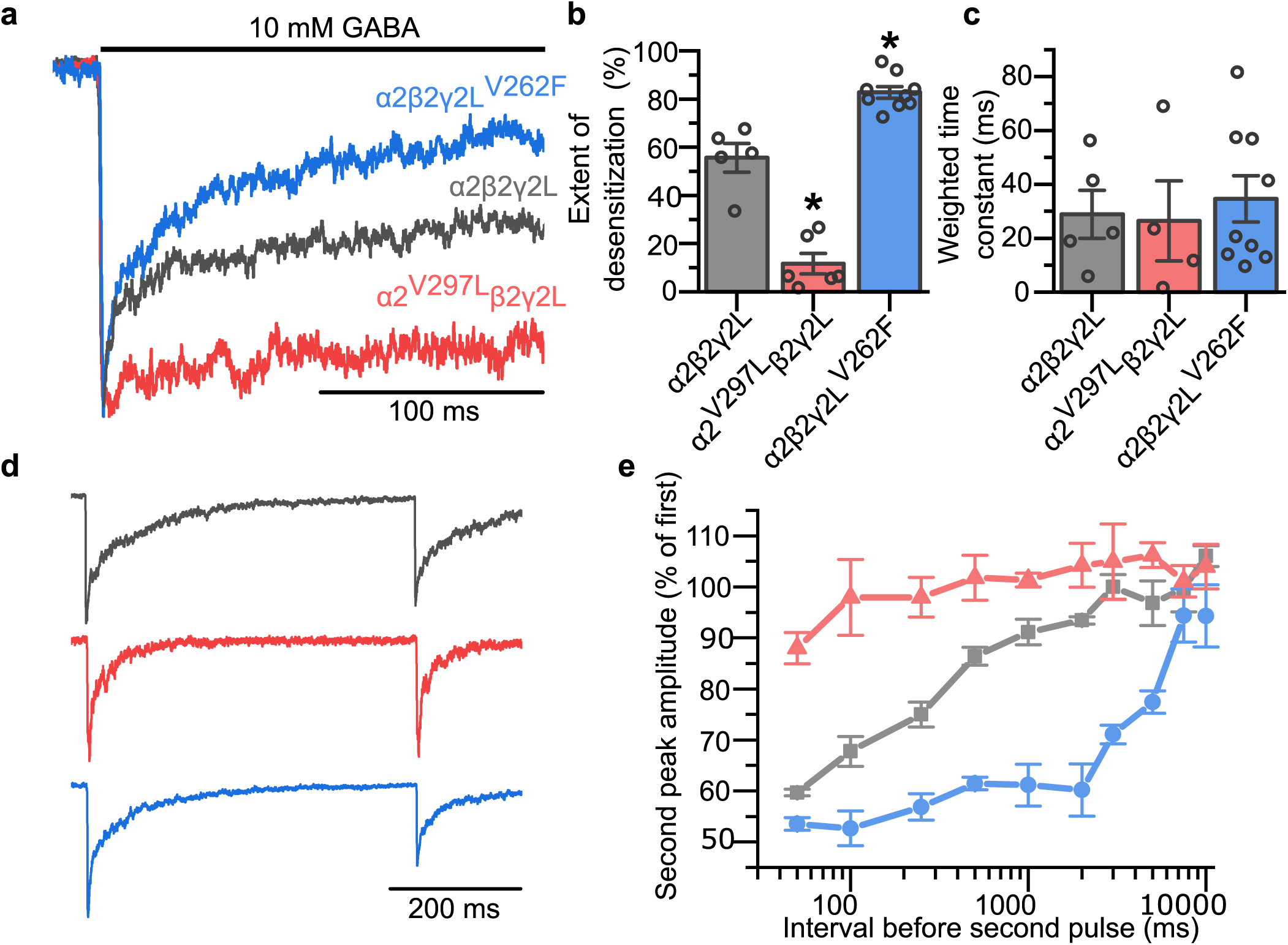
α2^V297L^ and γ2^V262F^ alter the desensitization of GABA_A_Rs. **a** Average GABA (10 mM)-activated currents showing macroscopic desensitization of wild-type and mutant α2 and γ2L GABA_A_Rs in outside-out patches pulled from transiently-transfected HEK293 cells expressing α2β2γ2L wild-type and mutated subunits, as indicated. **b** Extents of desensitization caused by 10 mM GABA measured as the % depression of the peak to steady-state GABA current, n = 5, 6, and 9 patches for receptors containing α2^WT^, α2^V297L^ and γ2L^V262F^, respectively. **c** Weighted decay time constants for exponential fits to macroscopic desensitization curves for α2^WT^, α2^V297L^ and γ2L^V262F^ containing GABA_A_Rs. **d** Averaged GABA currents recorded from outside-put HEK cell patches expressing GABA_A_Rs (colour coded as in **a**) showing responses to two 1 ms pulses of 10 mM GABA separated by 500 ms. **e** Re-sensitization of the peak GABA current measured from the relative amplitude of the current evoked by the second GABA pulse over a series of inter-event intervals, n = 4 patches for each.

To understand how these mutations might affect synaptic inhibition, a paired-pulse protocol was implemented, with outside-out patches exposed to two 1 ms pulses of 10 mM GABA separated by varying intervals between the pulses to monitor the recovery from desensitization. For α2β2γ2L wild-type receptors, the peak of the second response was depressed compared to the first response and this depression gradually recovered as the interval between GABA applications was extended over a period of 2-3 seconds (Fig. 2d,e). However, the inclusion of the α2^V297L^ subunit largely abolished the suppression of the second pulse for all inter-event intervals (Fig. 2d,e). By contrast, whilst γ2L^V262F^ only slightly increased the depression of the second current for short inter-event intervals, it substantially prolonged the duration over which the suppression was observed compared to wild-type (Fig. 2d,e), an observation entirely consistent with the stabilisation of the desensitized state.

To assess how manipulating the ability of GABA_A_Rs to desensitize might alter the strength of inhibitory synapses, the mutants were expressed in hippocampal neurons in culture. At room temperature, the expression of either the wild-type α2 or γ2L subunits, or the α2^V297L^ mutation did not alter the basal amplitudes of sIPSCs compared with mock-transfected control neurons (Fig. 3a-c; one-way ANOVA: F_(4, 57)_ = 5.68, p = 0.00064; Tukey test (mock, α2^V297L^): p = 0.99). Conversely, expressing the profoundly desensitising γ2L^V262F^ mutant substantially increased sIPSC amplitudes from −65.8 ± 9.4 pA (mock transfection control) to −126.7 ± 12.0 pA and also their rate of rise (Fig. 3a-c, g, h); Tukey test (mock, γ2l^V262F^): p = 0.0022). This was contrary to our initial expectation, based on the effects of this mutation on receptor kinetics, since the enhanced paired-pulse suppression exhibited by this mutant might be expected to manifest as a decrease in mean sIPSC amplitudes. However, the frequency of the synaptic events observed in these cultures were likely too low to observe a similar paired-pulse suppression, and indeed, the amplitudes of the sIPSCs did not show any negative correlation with their frequencies (Fig. 3d). Additionally, no significant changes were observed in either the interevent intervals (Fig. 3f; one-way ANOVA: F_(4,57)_ = 1.72, p = 0.16); or the 20-80 % rise times of the events, other than a small difference between the mutants (Fig.3e; one-way ANOVA: F_(4,57)_ = 3.24, p = 0.017; Tukey test (V297L, V262F): p = 0.013).

**Figure 3.**
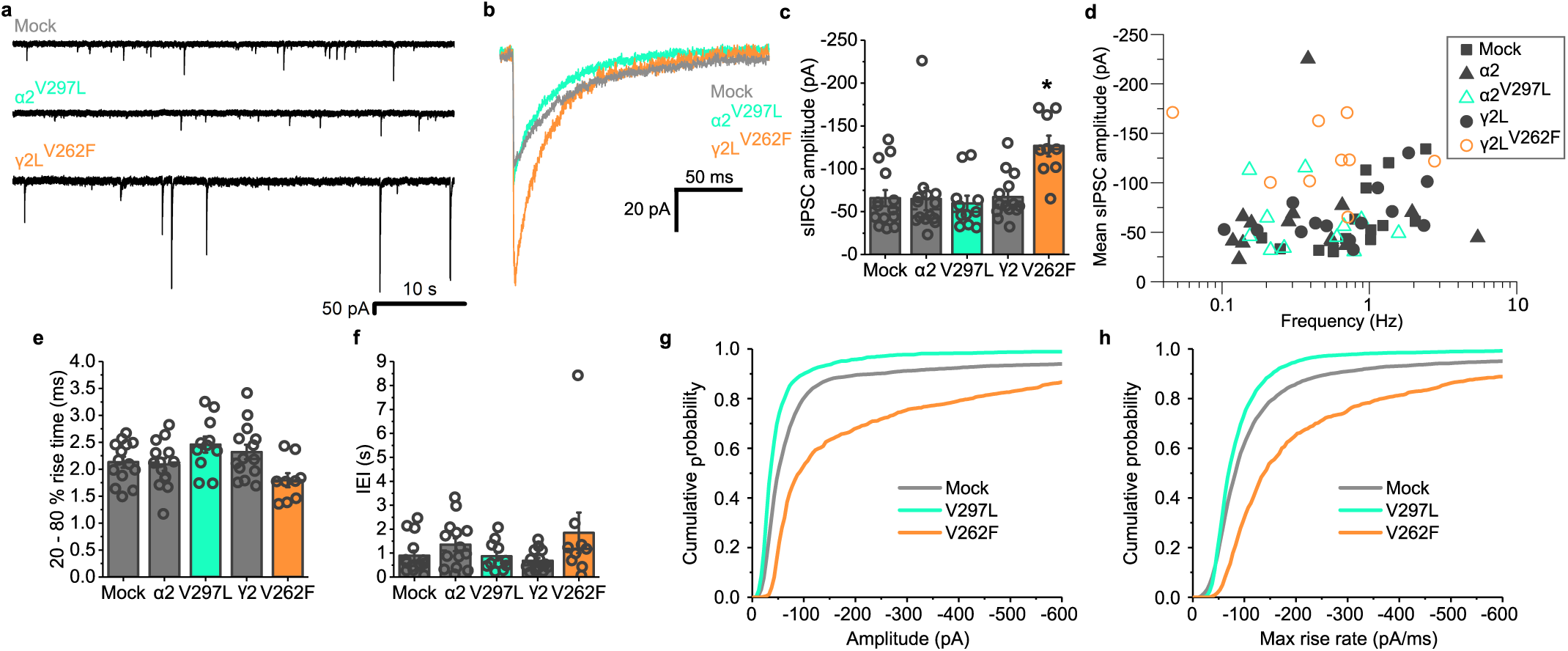
γ2^V262F^ expression increases sIPSC amplitudes at room temperature. **a** Representative sIPSC traces, recorded at 21°C from voltage clamped neurons expressing the indicated constructs. **b** Average waveforms of sIPSCs recorded for each receptor construct, aligned on the rising phase and not peak scaled. **c** Mean sIPSC amplitudes for neurons either mock-transfected, or expressing α2 or γ2L wild-type, or α2^V297L^, γ2L^V262F^; n = 14, 14, 11, 14, and 9 neurons, respectively. **d** Mean sIPSC amplitude plotted against mean sIPSC frequency for neurons transfected with the indicated constructs. **e** Mean 20-80% rise time of the recorded cells. **f** Median IEIs for all cells. **g** Mean cumulative probability distributions of the amplitudes, and **h** maximum rate of rise for the slopes of all sIPSCs recorded; n = 4335 events from 14 cells (Mock), 3971 events from 11 cells (a2^V297L^) and 1593 events from 9 cells (g2L^V262F^).

To assess the effects of the mutations under physiological conditions and increase the synaptic event frequency in the cultures, the experiment was repeated at 37°C. Here, similar results were observed, with γ2L^V262F^ increasing the average amplitudes of the synaptic events from −133.6 ± 8.9 pA (wild-type) to −211.1 ± 25.0 pA (Fig. 4a-d; one-way ANOVA: F_(4, 78)_ = 6.67, p = 0.00011; Tukey test (mock vs α2^V297L^): p = 0.96; Tukey test (mock vs γ2l^V262F^): p = 0.0022). Although the frequencies were higher under these conditions, again no correlation was evident with the sIPSC amplitudes (Fig. 4d). However, it is important to note that the events were selected for this analysis on the basis of having smooth uninterrupted rising phases lacking secondary peaks. This would have reduced the presence of summated sIPSCs in the dataset. To check for unforeseen consequences of our event preselection, we also examined the distributions of all detected events irrespective of their rising phase profiles (Fig. 4e-h). Interestingly, when all synaptic events are included in the analysis, the increase in sIPSC amplitude caused by γ2L^V262F^ was smaller (Fig. 4g; mean of mock transfection distribution = − 160.08 pA, and γ2L^V262F^ = −203.92 pA). This was to be expected due to the enhanced receptor desensitization conferred by this mutant, thereby reducing the ability of synaptic responses to summate in response to high probability GABA release at 37°C (e.g. Fig. 2d,e). Consistent with reduced summation, all the synaptic events recorded from cells expressing this mutant exhibited a decrease in their rise times (Fig. 4f; one-way ANOVA: F_(4,78)_ = 5.51, p = 0.00059 Tukey test (γ2^V262F^): p = 0.0024). Additionally, events were detected with a lower frequency for these cells (Fig. 4e; one-way ANOVA: F_(4,78)_ = 3.49, p = 0.011; Tukey test (mock, V262F): p = 0.0099).

**Figure 4.**
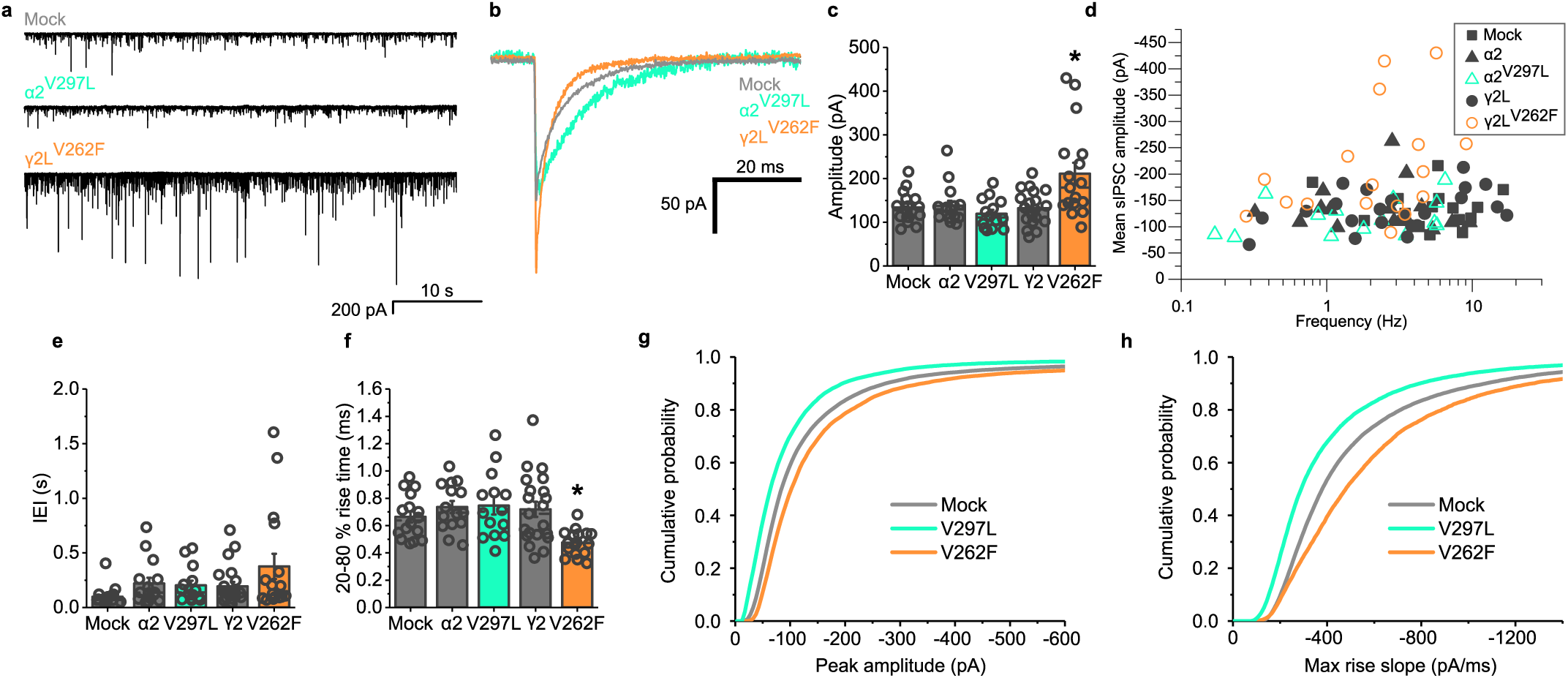
γ2^V262F^ expression increases sIPSC amplitudes at physiological temperature. **a** Representative sIPSCs recorded at 37°C from voltage clamped neurons expressing the indicated constructs. **b**, Average sIPSC unscaled waveforms and **c**, summary bar graph for mean sIPSC amplitudes: n = 16 (mock transfected), 15, 14, 21, and 17 neurons, for each construct as they appear on the abscissa. **d** Mean sIPSC amplitude plotted against average sIPSC frequency for each construct (see symbol key). **e** Median IEIs for all cells. **f** Mean 20-80% rise times for all cells. **g** Mean cumulative probability distributions of the amplitudes and **h** maximum rate of rise slopes of all sIPSCs recorded. n = 54092 events from 16 cells (Mock), 24377 events from 14 cells (a2^V297L^), and 24644 events from 17 cells (g2L^V262F^).

To further elucidate the cause of the increase in sIPSC amplitudes by V262F, the experiment was repeated in the presence of tetrodotoxin (TTX) to block action potential-dependent GABA release; however, the resulting miniature event (mIPSC) amplitudes were still enhanced by γ2L^V262F^ (Fig. 5a-d; t test vs mock: t_(9)_ = −2.87, p = 0.019). Additionally, the rise time of the mIPSCs for V262F-expressing cells was still faster compared to mock-transfected cells (Fig. 5e; t-test: t_(9)_ = 2.84, p = 0.019). However, the median synaptic event frequency was slower, suggesting a reduced release probability compared to mock-transfected (Fig. 5f; t-test: t_(9)_ = − 2.52, p = 0.033).

**Figure 5.**
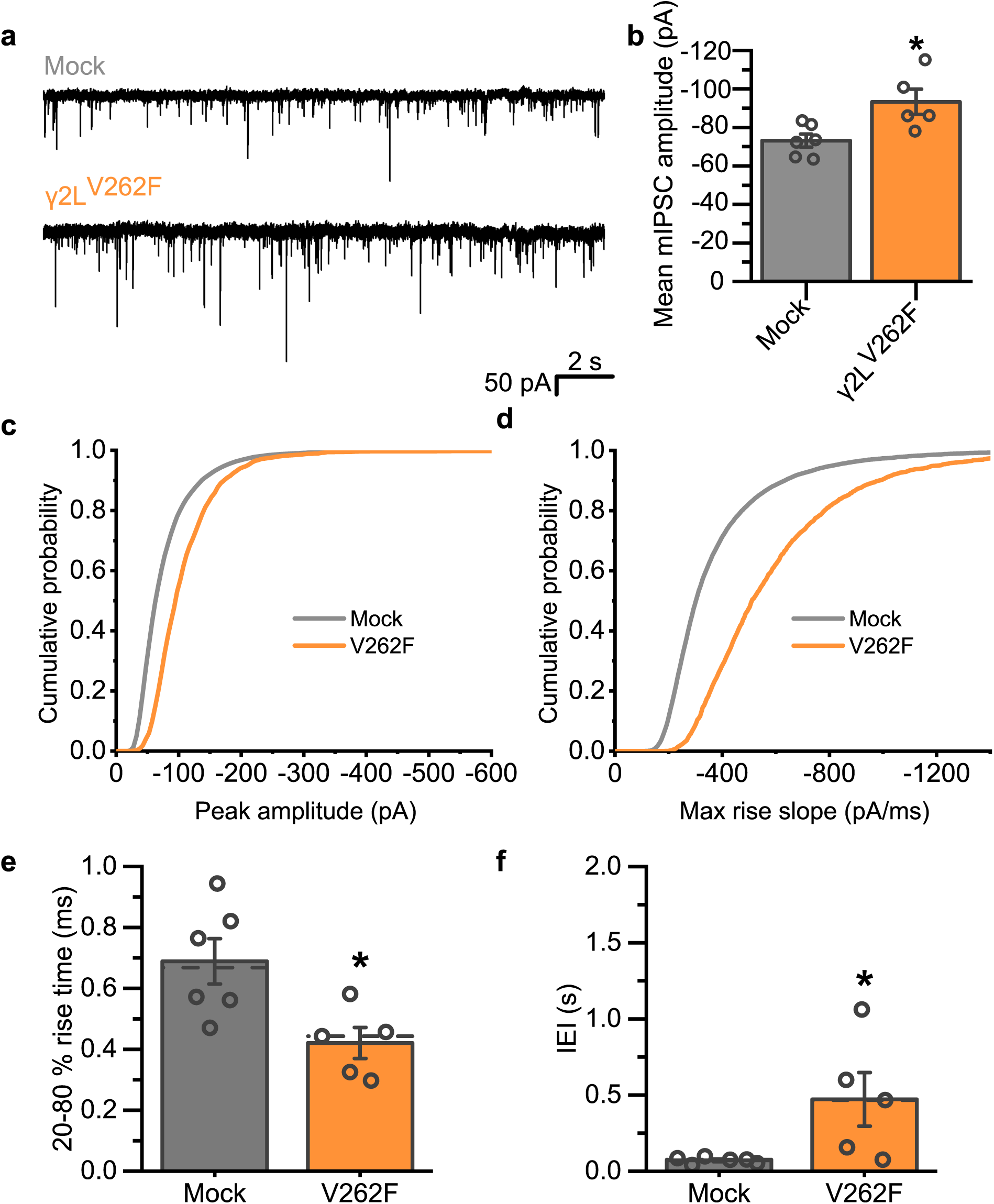
γ2^V262F^ expression increases miniature IPSC amplitudes. **a** Representative mIPSCs recorded in voltage clamp mode from neurons in the presence of tetrodotoxin (500 nM) either mock-transfected or expressing γ2^V262F^. **b** Mean amplitudes of mIPSCs for the constructs shown in **a. c** Cumulative probability distribution of mIPSC amplitudes and **d** maximum rate of rise slopes of all mIPSCs recorded; n = 18668 events from 6 cells (mock), and 3742 events from 5 cells (γ2L^V262F^). **e** Mean 20-80% rise times of the mIPSCs recorded from each cell. **f** Median IEIs for the mIPSCs recorded from each cell.

Given these observations, the profound increase in synaptic current amplitudes by γ2L^V262F^ is unlikely to be accounted for by an effect on GABA_A_R kinetics, as entry into the desensitized state alone would be expected to decrease IPSC amplitudes^16^. Moreover, it is also unlikely to be a consequence of an alteration in neural network activity given that it is still observed for both spontaneous and miniature inhibitory events. Instead, it is more likely that the expression of this mutant alters the numbers of GABA_A_Rs at inhibitory synapses, either through desensitization acting to regulate receptor number, or perhaps due to a homeostatic response to passing smaller Cl^-^ currents at synapses containing this mutant subunit during intense periods of synaptic activity.

### Agonist-induced LTP at inhibitory synapses depends on GABA_A_R desensitization

To examine whether the entry of receptors into the desensitized state is a necessary prerequisite for the induction of plasticity, the GABA pre-treatment protocol was repeated, but with the sIPSCs recorded earlier, approximately 20 min post-treatment, rather than 24 hr. Even at this early time-point, sIPSC amplitudes were substantially increased from −55.7 ± 3.8 pA (vehicle) to −132.2 ± 16.8 pA (post-GABA; Fig. 6a-d, f; one-way ANOVA: F_(3, 31)_ = 13.85, p = 0.0000066; Tukey test (vehicle vs GABA): p = 0.00002). By contrast, recordings from neurons expressing the minimally desensitising α2^V297L^ mutant revealed no potentiation of sIPSCs (Fig. 6c, d, f; Tukey test (vehicle, α2^V297L^ + GABA): p = 0.99). These results are consistent with the induction of agonist-induced iLTP requiring entry of GABA_A_Rs into a desensitized state. Indeed, α2^V297L^ should enhance the steady-state membrane Cl^-^ conductance by reducing the ability of the receptor to desensitize, yet this perturbation to Cl^-^ flux does not contribute to iLTP. Spontaneous IPSC decay kinetics (Fig. 6g; T test: t_(15)_ = −0.33, p = 0.74), event frequencies (Fig 6e; one-way ANOVA: F_(3, 31)_ = 1.33, p = 0.28), and rise times (Fig. 6h; one-way ANOVA: F_(3,31)_ = 0.97, p = 0.41) were unaffected by the pre-treatment with GABA, suggesting that the observed change in event amplitudes did not result from either changes in probability of release or receptor kinetics.

**Figure 6.**
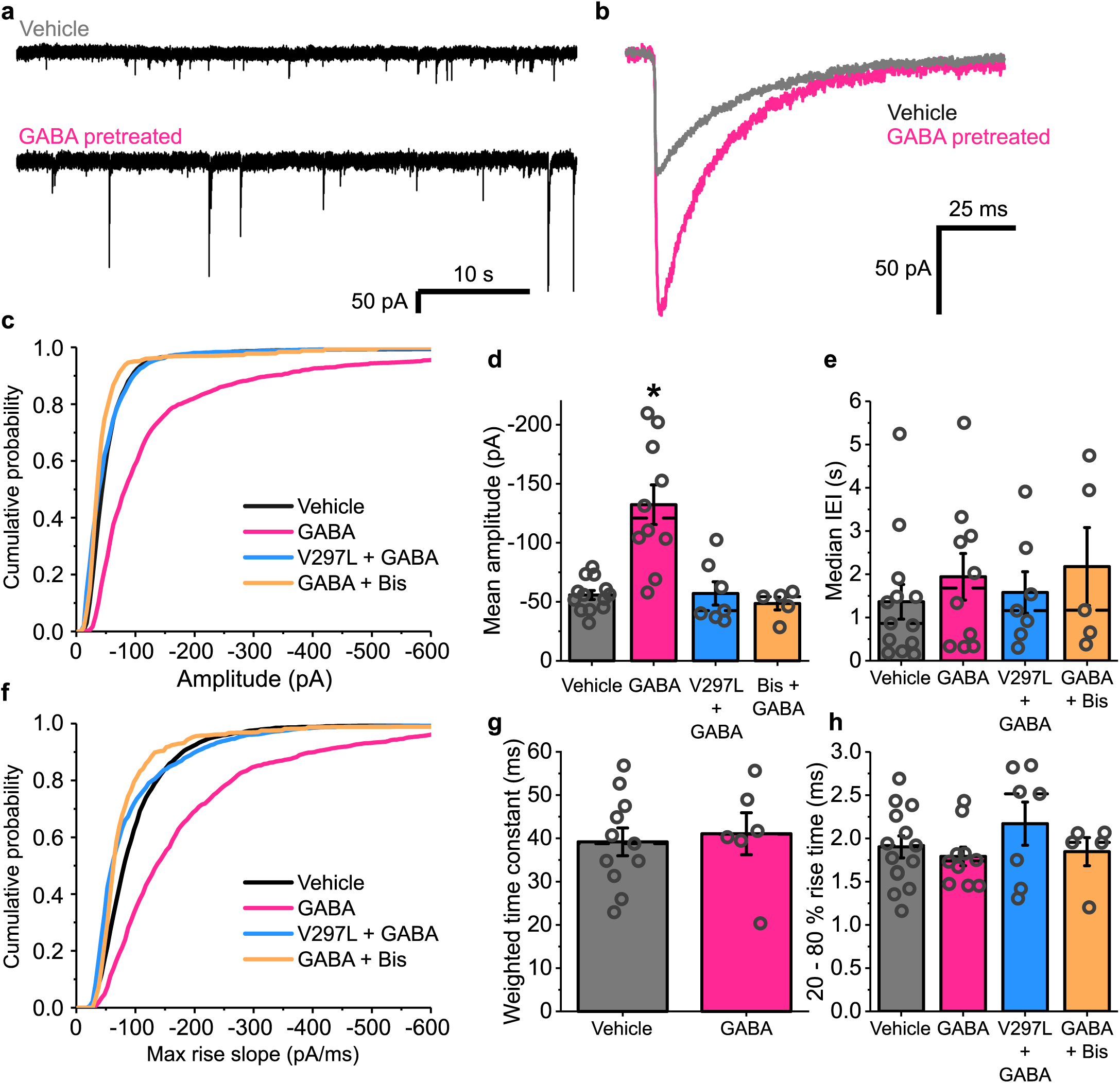
Induction of iLTP by GABA requires both GABA_A_R desensitization and PKC activation. **a** Representative sIPSC traces recorded in voltage clamp mode from neurons approximately 20 mins after treatment with either GABA (1 mM) or vehicle. **b** Average, unscaled sIPSC waveforms aligned on the rising phase. **c** Cumulative probability distribution of the amplitudes of all sIPSCs recorded: n = 4178 events from 13 cells, 2797 events from 10 cells, 1384 events from 7 cells, and 1440 events from 5 cells, respectively for each construct as they appear in the key. **d** Mean amplitudes of sIPSCs recorded from neurons pre-treated with either vehicle or GABA, pre-treated with GABA whilst expressing α2^V297L^, or pre-treated with GABA in the presence of bisindolylmaleimide-I (Bis). **e** Median IEIs for all events recorded from neurons after the indicated treatments. **f** maximum rate of rise slopes for all sIPSCs recorded. **g** Weighted time constants of exponential fits of the decay phases of sIPSCS recorded after treatment with either vehicle or GABA. **h** Mean 20-80 % rise times of all events recorded for each neuron in the indicated conditions.

Exposing neurons to GABA for prolonged periods has been reported to initiate a persistent increase in the PKC-dependent phosphorylation of the GABA_A_R γ2 subunit^21^. From prior studies^42^, we hypothesized that such a post-translational modification could, in principle, account for an increase in inhibitory synaptic efficacy, particularly since PKC phosphorylation of the γ2 subunit correlates with the clustering of GABA_A_Rs at inhibitory synapses^26,43,44^. To test this concept, GABA pre-treatment was repeated in the presence of the specific PKC inhibitor bisindolylmaleimide-I (Bis, 500 nM). This treatment completely blocked the induction of agonist-induced iLTP (Fig. 6c-f, h; Tukey test (vehicle vs GABA + bisindolylmaleimide): p = 0.97).

### Inhibitory LTP is independent of receptor kinetics and probability of GABA release

Given that the sIPSCs recorded here likely represent a mixture of responses to GABA release from different interneurons with different release probabilities, we explored whether the observed increase in sIPSC amplitude following GABA exposure was segregating with subsets of rise time and/or sIPSC frequency-related parameters. This enabled us to probe whether the potentiation was limited to specific subsets of events or instead represented an overall effect on inhibitory neurotransmission. To address this, we applied skewed t-distribution mixture modelling to a subset of sIPSC variables recorded for each synaptic event. Variables were selected to represent the rise time and frequency characteristics of the events but were independent from the event amplitude (see Methods). In adopting such criteria, our aim was to avoid simply isolating clusters of events that have undergone amplitude potentiation and to only reveal clusters with our selected parameters. When viewed using t-distributed stochastic neighbour embedding (t-SNE) or as a plot of rise time against frequency, these parameters were observed to have complex, overlapping, distributions (Fig 7a,b). Nevertheless, this analysis strategy yielded a separation of the data into 10 discernible clusters (Fig. 7a-c). Examination of the properties of each cluster revealed that events were segregated by instantaneous frequency (Fig. 7b,f), along with a separation between events exhibiting fast- and slow-rise times (Fig. 7b,e). Notably, despite the number of clusters, the amplitude potentiation of events following pre-exposure to GABA was relatively evenly distributed amongst all 10 clusters, (for both fast and slow rising events), with the spectrum of event frequencies demonstrating potentiation as increments in both peak amplitude (Fig. 7d), and the slopes of their rises (Fig. 7g,h) following exposure to GABA for 20 min at 1 mM.

**Figure 7.**
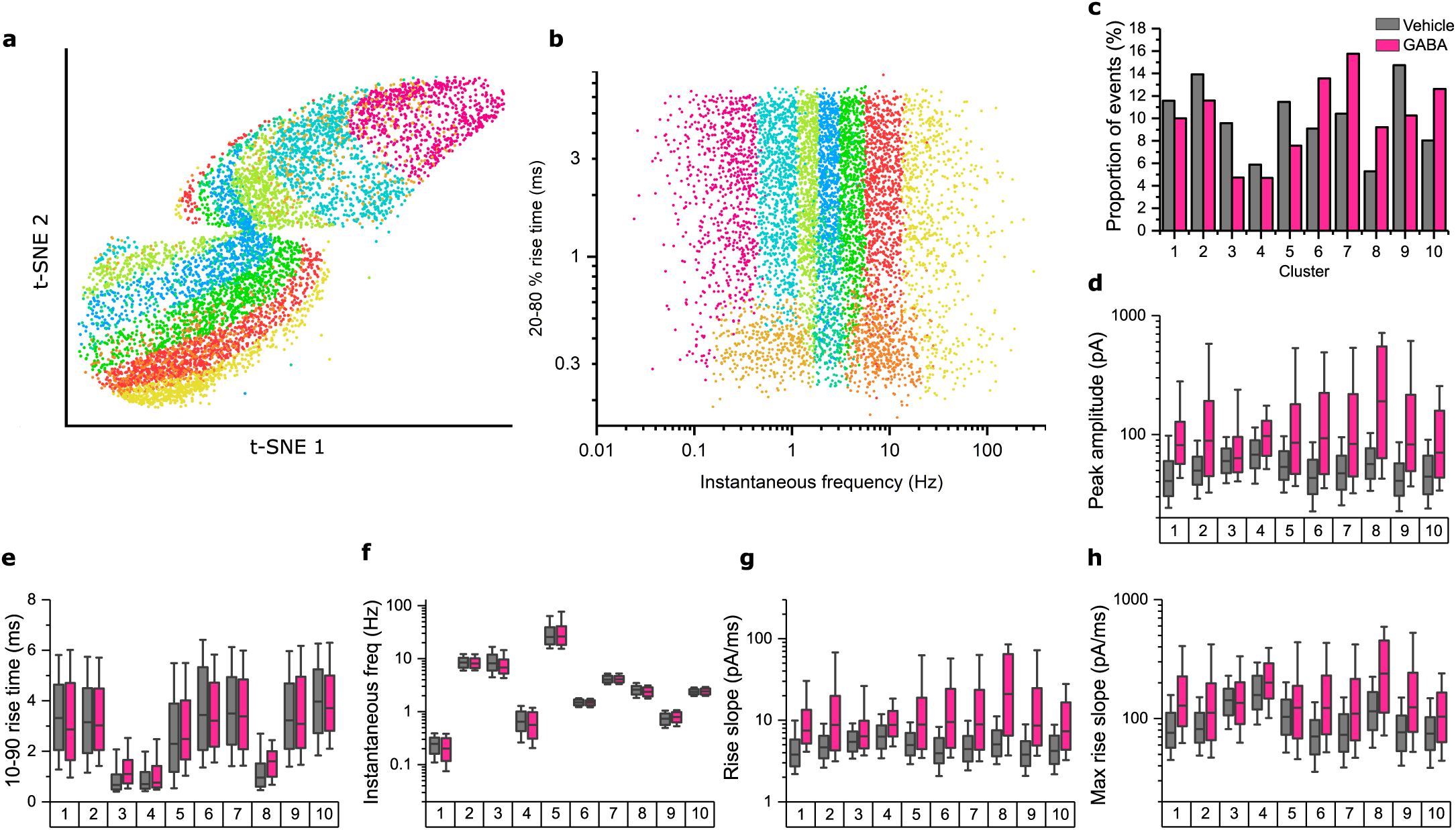
Cluster analysis of sIPSCs. **a** t-distributed stochastic neighbour embedding (t-SNE) plot of all mock/GABA-pretreated sIPSCs from figure 6. The t-SNE plot was produced using the same parameters as were used to analyse the clustering of events (see Methods). Events are colour coded according to the clustering determined by skewed mixture modelling of the data (10 total clusters). **b** Plot of 20-80 % rise time against instantaneous frequency of all sIPSCs, coloured by cluster. **c** Proportion of vehicle and 1 mM GABA-pretreated IPSC events in each cluster identified by the skewed mixture modelling. **d-h** Comparison of vehicle against GABA-treated events for each of the 10 clusters, for: **d** peak IPSC amplitude, **e** 10-90 % rise time, **f** instantaneous frequency, **g** rise slope, and **h** maximum rise slope.

### Direction of GABA Cl^-^ flux and iLTP

Some previous studies of agonist-dependent plasticity at inhibitory synapses have suggested the involvement of underlying mechanisms that require depolarising currents mediated by GABA_A_Rs to induce Ca^2+^ signalling^29,45^. These studies have associated depolarising GABA-currents with reduced synaptic efficacy. Although the evidence from our desensitisation mutants suggests that iLTP induction is not consistent with the involvement of any change in Cl^-^ current, we nevertheless assessed whether the GABA currents induced during the pre-treatment protocol are inhibitory or excitatory in nature.

To do so, we recorded from neurons under near-identical conditions to those during the GABA pre-treatment (i.e. bathed in culture media, with 5% CO_2_ / 95% O_2_, at 37°C). We used the cell-attached patch configuration to avoid perturbing Cl^-^ homeostasis. Under these conditions, the neurons exhibited robust spike firing. However, GABA (1 mM) application immediately silenced firing (Fig 8a, b). Conversely, application of bicuculline (50 μM) increased spike frequency (Fig 8c, d). These results are consistent with GABA_A_R mediated Cl^-^ currents in our cultures causing inhibition, and so discount depolarising GABA effects as the cause of the observed plasticity.

**Figure 8.**
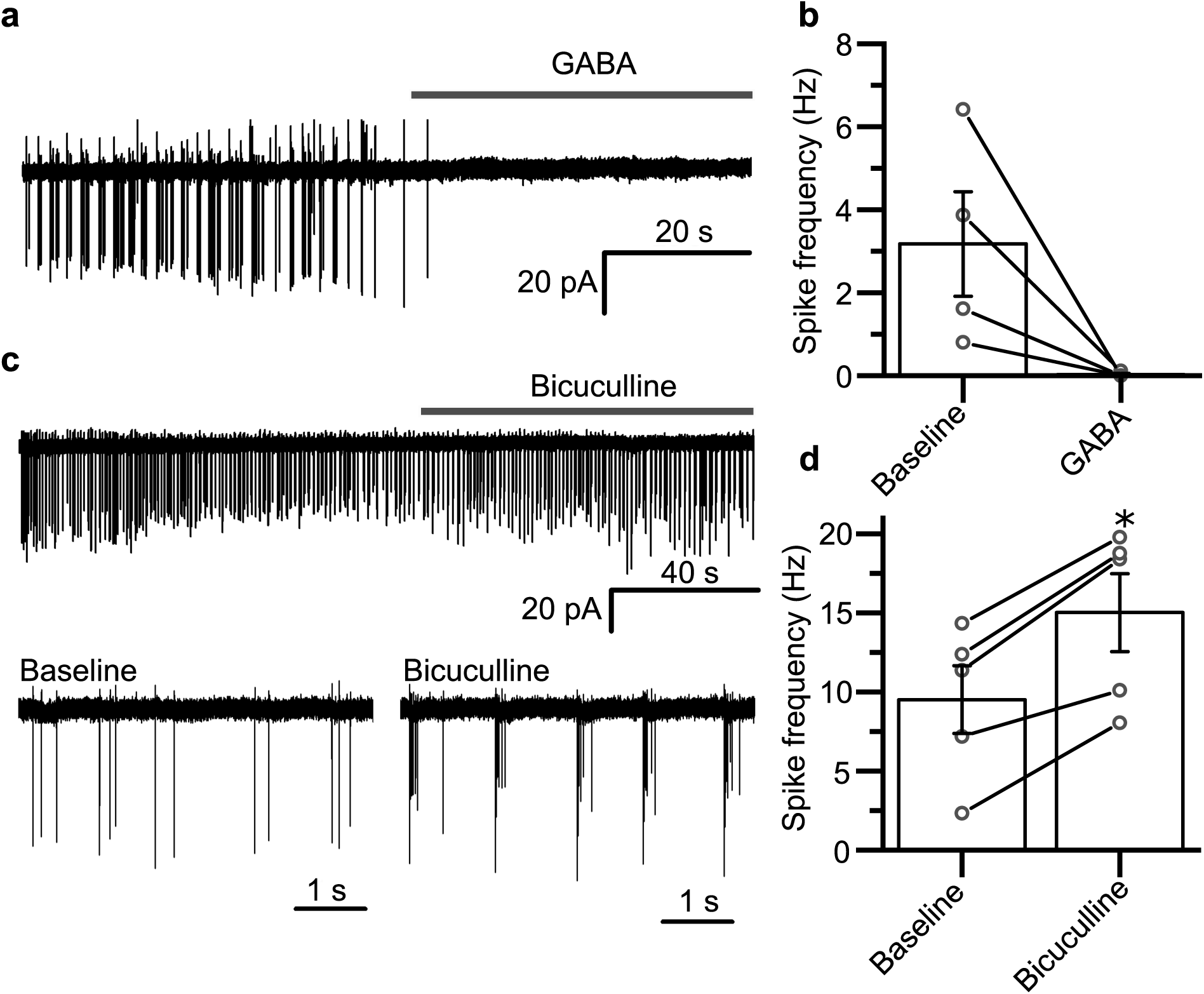
GABA is inhibitory under conditions that induce iLTP. **a** Extracellular spike currents recorded from a cultured hippocampal neuron in cell-attached voltage-clamp mode showing the response to 1 mM GABA application. **b** Summary of spike current frequencies before and after GABA exposure, n = 4 neurons. **c** Representative current trace recorded from a cultured hippocampal neuron in cell-attached voltage-clamp mode showing the response to bicuculline (50 µM) application – two segments of the recording are also shown below with higher time resolution. **d** Summary of spike current frequencies before and after bicuculline application, n = 5 neurons. Baseline measures signify the spike firing rate before any treatment.

### GABA_A_R desensitization increases phosphorylation of γ2^S327^ to induce iLTP

We next considered what could be the most plausible mechanism for the induction of iLTP. It is highly likely that receptor phosphorylation is key, evident from our observation that the induction of agonist-dependent iLTP requires PKC activity, combined with previous reports suggesting that such treatment also increases phosphorylation of the γ2 subunit^21^. Specifically, we examined whether GABA_A_R desensitization could increase the phosphorylation of key residues on γ2 subunits that are known to be associated with receptor clustering^26,43^. This could, in principle, lead to an increase in the number of receptors clustered at inhibitory synapses which would, in turn, account for the increase in sIPSC amplitudes.

To determine whether a change in the number of synaptic receptors is responsible for the potentiated sIPSCs and for iLTP, peak-scaled non-stationary variance analysis was performed on sIPSCs recorded from GABA pre-treated neurons (Fig 9a). Parabolic curve fits to the sIPSC variance-amplitude relationships for each cell showed no alteration in the unitary current (I_u_) passed by each receptor, when compared to those for vehicle-treated cells (Fig 9a-b; −3.33 pA for vehicle, −2.86 pA for GABA pre-treated; t_(21)_ = −1.17, p = 0.25), discounting any changes in single channel conductance or Cl^-^ equilibrium potential (under these conditions) as being causative of iLTP. However, the same curve fits revealed large increases in receptor numbers (N) activated at the peak of the sIPSC, more than doubling from 25.9 (mock) to 63.3 (GABA pre-treated) (Fig 9a, c; t_(21)_ = −4.48, p = 0.00021). This is consistent with an increase in the number of receptors mediating synaptic responses, sufficient to account for the increase in average sIPSC amplitude (Fig 6d).

**Figure 9.**
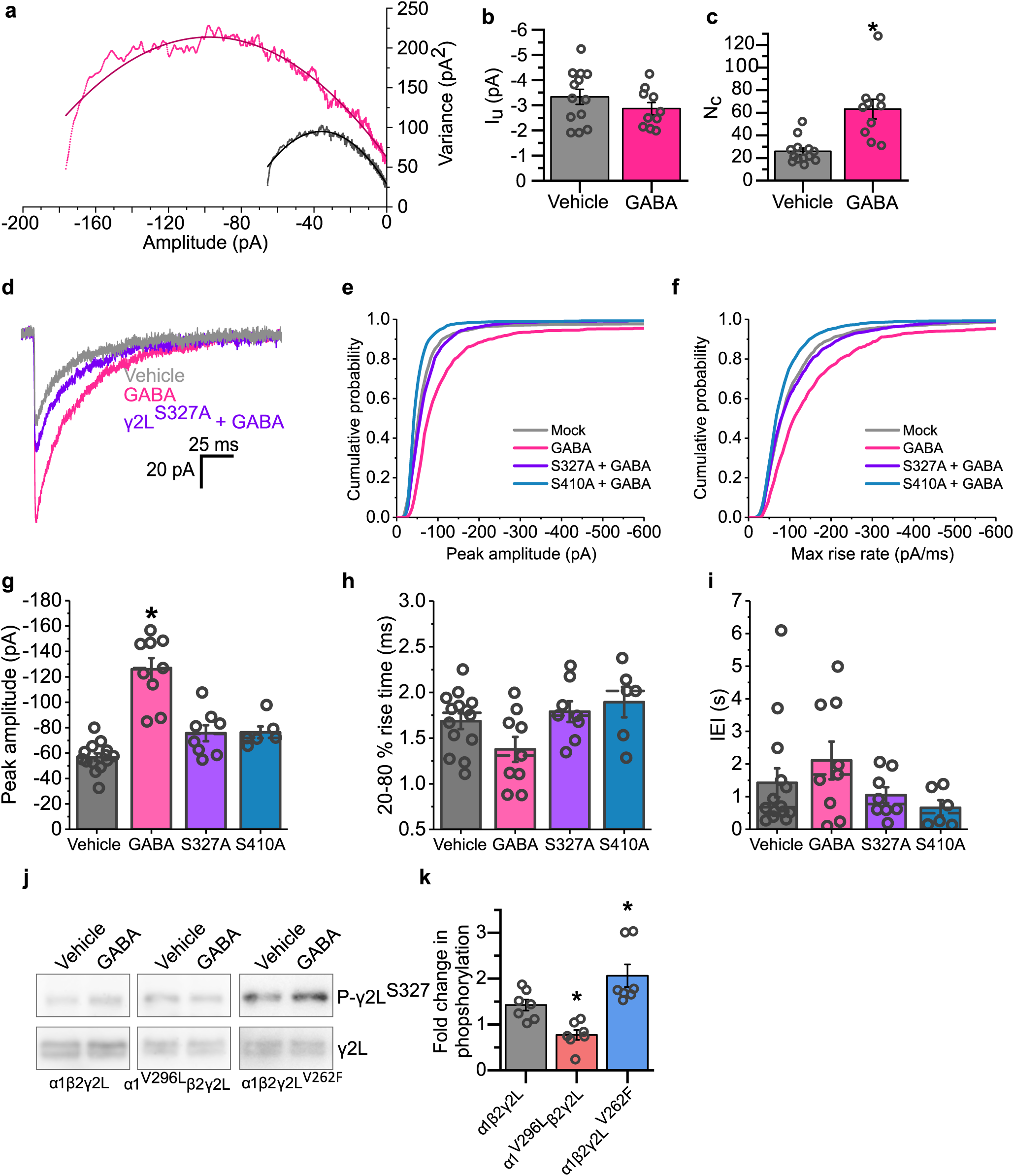
GABA_A_R desensitization increases synaptic receptor numbers by phosphorylating γ2^S327^. **a** Plots of synaptic current amplitude variance against current amplitude from peak-scaled non-stationary variance analysis of sIPSCs presented in fig 6 (n = 13 and 10 cells respectively). Data are shown for sIPSCs recorded from all GABA pre-treated neurons (pink) and from controls (black). **b** Single channel current amplitudes (I_u_) are calculated for synaptic GABA_A_Rs for each treatment. **c** Synaptic receptor numbers (N_c_) calculated for sIPSCs for each treatment. **d** Averaged sIPSC waveforms recorded from untransfected cells pre-treated with either vehicle or GABA, and cells expressing γ2^S327A^ pre-treated with GABA, aligned to their rise phases. **e** Mean cumulative probability distributions of the sIPSC amplitudes and **f** maximum rate of rise slopes of all sIPSCs recorded; n = 3470 events from 14 cells, 3808 events from 9 cells, 2256 events from 8 cells, and 4123 events from 5 cells respectively for each construct as they appear in the key. **g** Mean sIPSC amplitudes for the same conditions as in panel d. **h** Mean 20-80 % rise times for sIPSCs recorded under the indicated conditions. **i** Median IEIs of the sIPSCs recorded from cells under the indicated conditions. **j** Immunoblots of lysates taken from HEK293 cells transiently transfected with the indicated constructs treated with either vehicle or GABA (1 mM) for 20 min prior to lysis. Blots were probed for phosphorylated γ2^S327^ (top row), and for total γ2 levels (bottom row). **k** Fold changes in the phosphorylation levels for γ2^S327^ in cells expressing the indicated constructs, after GABA 20 min treatment; n = 7 lysates for each condition.

To assess whether this apparent increase in receptor number is indeed dependent on the phosphorylation of the γ2 subunit, we first repeated the GABA pre-treatment protocol on neurons expressing the γ2L^S327A^ construct which removes a key phosphorylation site. Again, pre-treatment with GABA increased the amplitudes of sIPSCs recorded from mock-transfected neurons, but this increase was largely abolished by the presence of the S327A phospho-mutant (Fig. 9d-g; one-way ANOVA: F_(3,33)_ = 29.20, p = 2.1×10^−9^; Tukey test (vehicle, S327A + GABA): p = 0.09), suggesting that the phosphorylation of this residue is necessary for the induction of agonist-induced iLTP. We also examined the involvement of an additional PKC-phosphorylation site on the β2-subunit (S410) by attempting to induce iLTP on neurons expressing the corresponding phosphorylation site mutation (S410A). Similar to S327, S410 also caused a complete absence of iLTP when exposed to GABA (Fig. 9e-g; Tukey test (vehicle vs S410A): p = 0.12), suggesting that iLTP may require the concerted action of PKC at multiple phosphorylation sites on the GABA_A_Rs. No alterations were observed in the frequencies or rise-times of the events under these conditions (Fig. 9h,i)

To directly probe whether desensitization can regulate the phosphorylation of S327 in γ2 subunits, we expressed α1β2γ2L, and the desensitisation mutants α1^V296L^β2γ2L, or α1β2γ2L^V262F^ GABA_A_Rs in HEK cells and assessed whether GABA treatments altered the level of its phosphorylation. Although phosphorylation of GABA_A_R subunits in response to prolonged agonist exposure has not previously been reported in HEK cells, phosphorylation has been observed directly after exposure to neurosteroids^18,19^. Indeed, we found that cells treated with GABA (1 mM) for 20 min increased the level of phosphorylation of γ2^S327^ by 42.3 ± 11.9 % compared to vehicle-treated controls for the wildtype α1β2γ2L receptor (Fig. 9j, k; one-sample t-test with a 1-fold change as the null hypothesis: t_(6)_ = 3.54, p = 0.013), similar to that previously reported for cultured cortical neurons^21^. Interestingly, this effect was completely blocked by expressing the non-desensitising α1^V296L^ mutant, for which a small decrease in phosphorylation relative to wild-type was also observed (77.2 ± 10.9 % phosphorylation in treated cells compared to vehicle; Fig. 9j, k: one-way ANOVA: F_(2, 18)_ = 14.19, p = 0.00020; Tukey test (wt vs α1^V296L^): p = 0.038). Conversely, GABA_A_Rs incorporating the profoundly desensitising γ2^V262F^ subunit displayed larger increases in phosphorylation of S327, relative to wild-type α1β2γ2L receptors, when exposed to GABA treatment (206.4 ± 24.9 %, Fig. 9j, k; Tukey test (wt vs γ2L^V262F^): p = 0.042).

The strong correlation between entry of the receptor into the desensitised state and the extent of γ2L^S327^ phosphorylation, and in particular, the observation that the increased phosphorylation was completely blocked by expression of the non-desensitizing α1^V296L^ mutant, strongly suggests that entry of the receptors into the desensitized state is required for agonist-induced γ2L^S327^ phosphorylation. This plausibly explains the dual dependence of the agonist-induced plasticity on the receptor’s desensitized and phosphorylation states.

### Desensitization facilitates phosphorylation of GABA_A_R subunits by activated PKC

The ability of desensitization to enhance the phosphorylation of GABA_A_R γ2L subunits by PKC could proceed via two principal mechanisms. Firstly, the desensitization of GABA_A_Rs could lead to the activation of PKC. Recent evidence indicates that GABA_A_R subunits can interact with and also activate phospholipase C (PLC)^26^. Although this results in the activation of calcineurin, with subsequent dephosphorylation of γ2L^S327^ and downregulation of inhibitory synapses, in principle this signalling pathway could also activate PKC and upregulate the receptors under different cellular signalling conditions. Similarly, it has also been reported that the desensitization of nicotinic acetylcholine receptors can activate G-protein mediated signalling^46^, suggesting that desensitization of Pro-loop receptors is indeed capable of initiating metabotropic signalling cascades.

The second mechanism by which GABA_A_R desensitization could increase phosphorylation of GABA_A_R subunits would be to facilitate the access of an already-active protein kinase to the substrate γ2L^S327^ residue. Consistent with such a mechanism, neurosteroids have previously been reported to increase the phosphorylation of GABA_A_R subunits without altering the phosphorylation of PKC itself^18^. Such a mechanism would presumably require conformational changes to the receptor that favour the presentation of the receptor substrate site(s) to the kinase.

Given that these two distinct types of mechanism would have different implications for the regulation of inhibitory synaptic plasticity, and therefore the context in which it might occur, an assay was devised to determine which form of regulation is more likely. This involved activating PKC in hippocampal neurons with phorbol 12-myristate 13-acetate (PMA, 200 nM). This treatment increased sIPSC amplitudes from −60.3 ± 8.3 pA to −104.8 ± 12.5 pA (Fig. 10a– e; one-way ANOVA: F_(3,25)_ = 5.84, p = 0.0036; Tukey test (vehicle vs PMA): p = 0.022). However, by co-incubating neurons with PMA and picrotoxin (Ptx, 100 μM) used to block GABA_A_R currents and desensitization during the pre-treatment phase, no increase in inhibitory synaptic current amplitudes was observed (Fig. 10c-e; Tukey test (vehicle vs PMA + Ptx): p = 0.98). Similarly, PMA pre-treatment of neurons expressing non-desensitising α2^V297L^ did not elicit any increase in sIPSC amplitudes (Fig. 10c-e; Tukey test (vehicle, α2^V297L^ + PMA): p = 0.99). These results suggest that even when PKC is activated, it is still necessary for GABA_A_Rs to undergo desensitization for the induction of iLTP. This scenario is more consistent with the second of our proposed mechanisms, as the first mechanism predicts that receptor activation and desensitization are superfluous if PKC is already activated.

**Figure 10.**
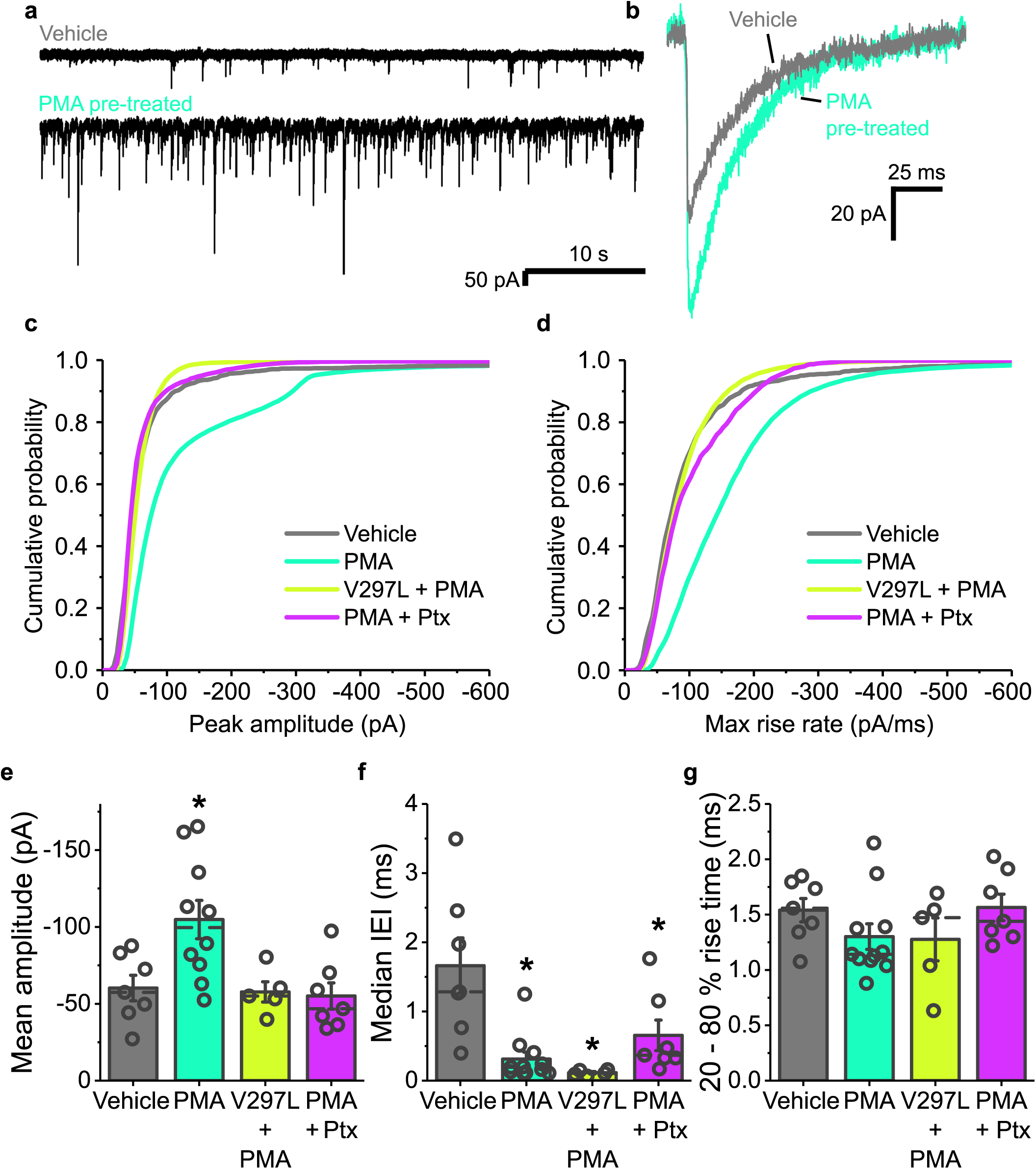
Induction of iLTP by PKC requires GABA_A_R activation and desensitization. **a** Representative sIPSC traces recorded from cells pre-treated with either vehicle or phorbol 12-myristate 13-acetate (PMA). **b** Average waveforms of sIPSCs for each condition. **c** Mean cumulative probability distributions of the amplitudes of all sIPSCs recorded for each receptor construct (WT unless specified otherwise) and condition as they appear in the key (Ptx = picrotoxin). **d** Cumulative probability distributions of the maximum rate of rise slopes of all sIPSCs recorded; n = 1482 events from 7 neurons, 12082 events from 10 neurons, 11168 events from 5 neurons, and 4757 events from 7 neurons respectively in the order they appear in the key. **e** Summary data for mean sIPSC amplitudes recorded from cells pre-treated with either vehicle or PMA, or cells expressing α2^V297L^ and pre-treated with PMA, or pre-treated with PMA in the presence of picrotoxin (100 µM). **f** Median sIPSC IEIs under the same four conditions as in e. **g** Mean 20-80 % rise time of sIPSCs for neurons recorded under the indicated conditions.

PMA also caused an increase in sIPSC frequency from 0.2 ± 0.06 Hz to 2.9 ± 0.6 Hz (Fig. 10f; one-way ANOVA: F_(3,25)_ = 8.55, p = 0.0004; Tukey test (vehicle vs PMA): p = 0.0007), and likely represents a distinct presynaptic action of PKC on GABA release as reported previously^47,48^. This decrease in IEIs was not blocked by expression of α2^V297L^ (Fig. 10f; Tukey test (vehicle vs α2^V297L^ + PMA): p = 0.0012). This would suggest that the increase in amplitudes caused by PMA was not simply a function of the increased release observed with this treatment. Consistent with this, no changes to the rise times of the events were observed under any of these conditions (Fig. 10g; one-way ANOVA: F_(3,26)_ = 1.33, p = 0.28).

### Allosteric modulators can induce desensitization-dependent LTP

Although entry of GABA_A_Rs into the desensitized state is commonly studied in the context of orthosteric agonists, allosteric modulators will also affect the occupancy of this state during synaptic inhibition. In the case of one negative allosteric modulator, pregnenolone sulfate, inhibition of GABA_A_Rs is caused by directly acting on the desensitized state, enabling either stabilisation or increased entry of GABA_A_Rs into this state^49-52^. However, given that entry into the desensitized state is thought to follow receptor activation^10,53^, it is also likely that drugs which do not act directly on the desensitized state will also alter its occupancy indirectly, as a consequence of effects on receptor kinetics. Following this line of reasoning, positive allosteric modulators should also increase the occupancy of the desensitized state.

To examine this hypothesis, we returned to the paired-pulse assay to assess GABA_A_R desensitization in outside-out patches in the presence of either pregnenolone sulfate, or the anaesthetic etomidate, which is a positive allosteric modulator of GABA_A_Rs^54^. Strikingly, both modulators had similar effects on the paired-pulse GABA current suppression (Fig. 11a, b) to that observed with the γ2L^V262F^ subunit mutant. This manifested as a suppression of the second GABA current at short inter-event intervals (similar to that for vehicle-treated receptors), but considerably more sustained at longer inter-event intervals in the presence of either modulator. This result is in accord with both modulators promoting the occupancy of agonist-bound closed states during phasic signalling, with etomidate also increasing the occupancy of open states evident from the slower decays of the evoked GABA currents (Fig. 11a, c).

**Figure 11.**
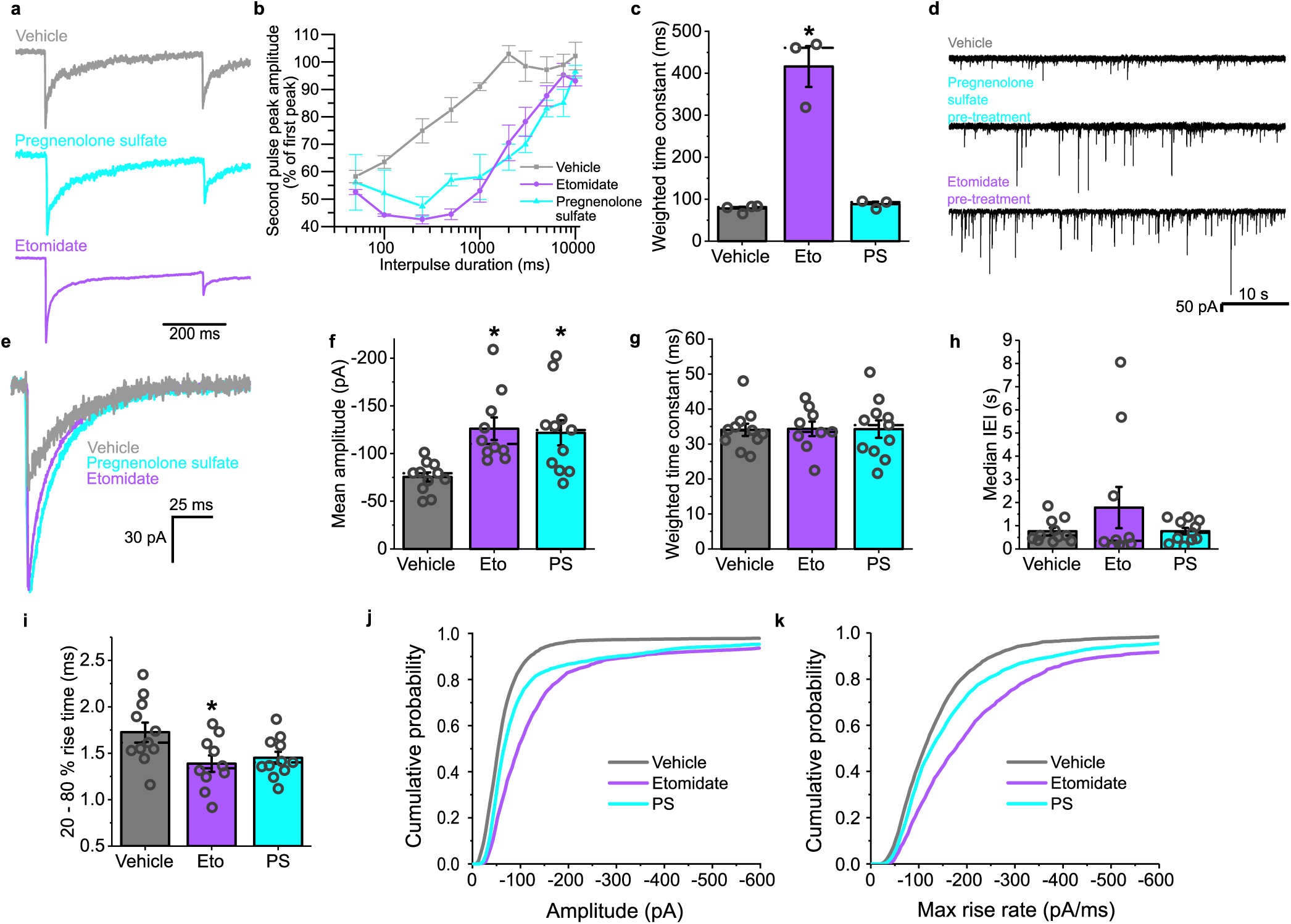
Allosteric modulators of GABA_A_Rs that increase the occupancy of the desensitized state also induce iLTP. **a** Averaged GABA currents of GABA_A_Rs in outside-out patches pulled from transiently transfected HEK293 cells, to sequential pulses of 10 mM GABA in the presence of vehicle, 10 μM pregnenolone sulfate (PS), or 5 μM etomidate. **b** Re-sensitization of the second GABA current with increasing inter-event intervals in the presence of the indicated modulators; n = 4, 3, and 3, respectively, for each condition (see key). **c** Weighted decay time constants for GABA current responses in control and exposed to the modulators (Eto = etomidate, PS = pregnenolone sulfate). **d** Representative sIPSCs recorded from neurons pre-treated with the indicated modulators (vehicle, 1 µM pregnenolone sulfate or 5 µM etomidate). **e** Average sIPSC waveforms for each modulator aligned on the current rising phases. **f** Mean sIPSC amplitudes and **g** weighted decay time constants for sIPSCs recorded after treatment with the indicated modulators. **h** Median IEIs for the sIPSCs recorded from neurons after treatment under the indicated conditions. **i** 20-80 % rise times of sIPSCs recorded after treatment. **j** Cumulative probability distribution of the sIPSC amplitudes. **k** Maximum rate of rise slopes of all sIPSCs recorded; n = 3929 events from 11 cells, 8638 events from 10 cells, and 5346 events from 11 cells, respectively in the order they appear in the key.

As both types of modulator appear to increase occupancy of the desensitized state during phasic signalling, we predicted that they would be capable of inducing iLTP. This is despite etomidate being a positive allosteric modulator of GABA_A_Rs and promoting activation and therefore GABA currents, and pregnenolone sulfate acting as a negative modulator to reduce GABA currents.

To test this hypothesis, we recorded from hippocampal neurons pre-treated with either modulator for 20 min prior to recording. For both types of modulator, this resulted in large increases in the amplitudes of sIPSCs, from −75.4 ± 4.8 pA (vehicle controls) to −121.8 ± 13.2 pA (1 µM pregnenolone sulfate) and −126.1 ± 11.8 pA (5 µM etomidate) (Fig. 11d-f; j-k; one-way ANOVA: F_(2, 29)_ = 7.19, p = 0.0029; Tukey test (mock vs PS): p = 0.0099; Tukey test (mock vs etomidate): p = 0.0058). The decay kinetics of the synaptic currents were unaltered in each of these recordings (Fig. 11e, g; one-way ANOVA: F_(2,28)_ = 0.0041, p = 0.99), consistent with each modulator having been washed out prior to recording. The event frequencies and rise times (Fig. 11h-i) showed no significant changes (one-way ANOVA (IEI) F_(2,29)_ = 4.25, p = 0.27; one-way ANOVA (rise times) F(2,29) = 4.25, p = 0.024), other than a slight decrease in the rise time of events recorded from neurons pre-treated with etomidate (Tukey test (Mock vs Etomidate): p = 0.029; Tukey test (Mock vs PS): p = 0.078).

Thus, the finding that either a positive or negative allosteric modulator can elicit prolonged increases in synaptic current amplitudes, subsequent to their washout, confirms that entry of the receptors into the desensitized state acts to induce iLTP at inhibitory synapses. Furthermore, this finding also suggests that this inhibitory plasticity can be induced using clinically relevant modulators of GABA_A_Rs.

## Discussion

In exploring and establishing the role that desensitization of GABA_A_Rs can play in the brain, we have uncovered a mechanism by which inhibitory synapses can directly regulate their level of plasticity, in the form of an inhibitory long-term potentiation that is manifest by increased synaptic current amplitudes.

In essence, iLTP is induced following the increased entry of GABA_A_Rs into one or more desensitized states, which appear to serve as a signal for the activity state of the inhibitory synapse. Receptor desensitization increases the phosphorylation of GABA_A_R subunits by PKC leading to a subsequent long-term increase in inhibitory synaptic efficacy. In this context, the physiological role of the desensitized state appears to be largely as a mechanism of signal transduction. This is a role to which it is well suited, especially given current structural and kinetic evidence^10,11,14,53,55^, as it represents a relatively long-lived conformational change transduced from the external to internal faces of the plasma membrane upon the binding of agonist.

This long-term form of inhibitory plasticity was induced with several chemical-LTP protocols stimulating either receptor desensitization (using GABA, pregnenolone sulfate or etomidate) or by direct PKC activation (using a phorbol ester). Similar increases in synaptic efficacy were also apparent in cells expressing a GABA_A_R mutation which substantially increases the stability of the desensitized state (γ2L^V262F^). The induction of iLTP could also be blocked by reducing entry into the desensitized state (using α2^V297L^), inhibition of PKC (using bisindolylmaleimide-I) or by preventing GABA_A_R activation and thus desensitization (with picrotoxin), or by ablating the sites of PKC phosphorylation on β2^S410A^ or γ2^S327A^.

In a physiological context, elevated levels of receptor desensitization will most likely occur during high intensity firing of interneurons, or in the presence of modulators such as endogenous neurosteroids or anaesthetics. The basal PKC activity required for this mechanism also has several potential sources in neurons, signalling through either brain-derived neurotrophic factor^56,57^, metabotropic glutamate receptors^58,59^ or other forms of neuromodulation^60,61^, all of which could potentially increase PKC activity leading to the phosphorylation of GABA_A_Rs and alterations in the efficacy of inhibitory synapses. Interestingly, a similar receptor-activity and PKC-dependent mechanism is already known to regulate the clustering of glycine receptors in the spinal cord^62^. Although the activity dependence was not demonstrated to be due to entry of the glycine receptors into their desensitized state, it was independent of the direction of net chloride ion flux. Furthermore, the structural mechanism of glycine receptor desensitization is similar to that of GABA_A_R^11,63^.

The phosphorylation of GABA_A_R subunits by PKC, following activation or modulation of the receptor, has been reported^18,19,21^. Additionally, although PKC activation is acknowledged to have variable effects on GABA_A_Rs^64^, the enhancement of inhibitory synaptic currents through PKC-dependent phosphorylation and the regulation of GABA_A_R clustering by the phosphorylation of the γ2^S327^ site are also well established^26,43,44,56,59,65^. Thus, the results described here provide a mechanistic basis for the link between GABA_A_R activation and phosphorylation, and demonstrate a synaptic relevance for this mechanism. Structural studies will be required to establish precisely how entry of the receptors into the desensitized state enhances the phosphorylation of GABA_A_R subunits in the M3-M4 internal loop domain, an area yet to be resolved at the atomic level.

Given that GABA_A_R activation and desensitization are prerequisites for iLTP even in the presence of PMA, it seems unlikely that this form of plasticity depends on the activation of phospholipase C by the GABA_A_Rs, even though such a signalling cascade has been described^26^, as PMA will occupy the diacylglycerol binding site of PKC^66^.

Interestingly, recent structural work has shown that phosphatidylinositol bisphosphate (PIP_2_) binds to GABA_A_Rs, with the inositol head group positioned in close proximity to the desensitization gate^67^. Moreover, the binding of lipids to this site has also been shown to decrease the desensitization of a Pro-loop receptor^68^. Thus, the entry of GABA_A_Rs into the desensitized state could favour the dissociation of bound PIP_2_, in the process presenting the inositol head group for binding to the C2-domain of certain isoforms of PKC. Consistent with this, recent structural work on the glycine receptor, and on the prokaryotic channel ELIC, has shown that the straightening of the M4 helix in a desensitized conformation disrupts this lipid binding site^69,70^. Furthermore, simulations of the analogous phosphatidylserine binding site on the glycine receptor have also shown that lipid binding is favoured in the inactive conformation relative to the active one^71^. Such a mechanism could effectively act as a signal of GABA_A_R activity at inhibitory synapses, potentially providing a more reliable indicator of synaptic activity than chloride fluxes or inhibition of excitatory signalling and thus enabling a greater range of inhibitory plasticity.

## Materials and Methods

### HEK cell culture

Human embryonic kidney cells (HEK293) were cultured in Dulbecco’s modified Eagle’s media (Gibco) containing 10 % v/v fetal bovine serum (heat-inactivated, Brazil, Gibco) and 100 u/ml penicillin and 100 μg/ml streptomycin (Gibco). They were maintained in a humidified 37°C, 5% CO_2_ incubator. Cells were passaged approximately twice weekly. Briefly, cells were detached from tissue culture plasticware using trypsin (Trypsin - EDTA, 0.05% w/v, Gibco). The dissociated cells were then centrifuged at 1000 rpm for 2 min before resuspension in culture media at the desired density for plating.

For electrophysiology experiments, cells were plated onto 22 mm washed glass coverslips coated with poly-L-lysine (mw: 70,000 – 150,000; Sigma). Cells were transfected with mouse GABA_A_R cDNA constructs using the pRK5 vector and with eGFP cDNA (pEGFP-C1) in a 1:1:1:1 ratio using the calcium phosphate precipitation method. For each coverslip 4 μg of total DNA was mixed with 20 μl of 340 mM CaCl_2_. This solution was then mixed with 24 μl 2x HBS (50 mM HEPES, 280 mM NaCl, 2.8 mM Na_2_HPO_4_). The combined solution was applied dropwise to the cells. Transfected cells were incubated in a humidified 37°C / 5% CO_2_ incubator for 24 - 48 hrs prior to recording. Cells that were incubated for 48 hrs were washed in Hanks balanced salt solution (HBSS, Sigma) after 24 hrs to remove the transfection media. For immunoblotting, a similar protocol was used with cells seeded onto 6-well plates.

### cDNA constructs

The constructs α1^V296L^ and γ2L^V262F^ were generated as described in Gielen et al (2015). The α2 equivalent of α1^V296L^ (α2^V297L^) was generated using site-directed mutagenesis of the wild-type α2 construct. Briefly, an inverse PCR was carried out with primers containing the mutation (forward:5’-CTGTTCTCTGCCCTAATTGAATTTGCA-3’;reverse:5’-AAACGCATAACAAACAGCTATAAACCAGTCC-3’). The products of the reaction were ligated with T4-ligase (Roche), and then introduced into competent *E. coli* cells (NEB 5αs, New England Biolabs). Clonal bacterial cultures were then produced, and the DNA harvested using a maxiprep kit (Qiagen, manufacturers protocol). All constructs were validated using Sanger sequencing (Source biosciences).

### Outside-out patch recordings from HEK293 cells

Outside-out patches were pulled from transfected HEK cells using thick-walled borosilicate patch electrodes (resistance 4 - 6 MΩ) filled with an internal recording solution consisting of: 140 mM KCl, 10 mM HEPES, 1 mM MgCl_2_, 4 mM Mg-ATP, 10 mM BAPTA, pH 7.2. The bath was continuously perfused with a Krebs solution containing: 140 mM NaCl, 4.7 mM KCl, 1.2 mM MgCl_2_, 2.5 mM CaCl_2_, 5 mM HEPES, 11 mM D-glucose, pH 7.4. The outside-out patches were voltage clamped at −20 mV with an Axopatch 200B amplifier (Molecular devices) and currents were recorded and low pass filtered with a 4-pole Bessel filter (5 kHz), at a sampling rate of 50 kHz. Drugs were rapidly-applied using a glass theta-tube, extruded to a tip diameter of 50 – 100 µm, and connected to a piezoelectric transducer (Burleigh Instruments) for fast drug exchange. Open electrode tip liquid junction currents were used to confirm the rate of solution exchange (10 - 90 %) at 100 – 250 µs, and that exposures were free from oscillation artefacts. All recordings from outside-out patches were made at 21 ± 1 °C.

GABA current decay curves were fitted using Clampfit version 10.7 (Molecular Devices) and the standard exponential function with two or three components. The number of components used was determined by whether additional components decreased the sum of squared errors. Weighted time constants (T_w_) were calculated with the formula:

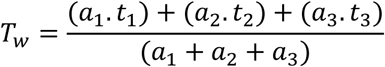

Where *a* indicates the amplitude, and *t* indicates the time constant, for each exponential component. Extents of desensitization were calculated as the percentage reduction in GABA current from the peak to the steady state:

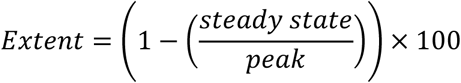

### Dissociated neuronal cultures

Hippocampal cultures were prepared from E18 Sprague-Dawley rat embryos. Animals were sacrificed in accordance with the UK Animals (Scientific Procedures) Act 1986. The pooled hippocampi were dissociated by trypsinization and trituration using flame polished Pasteur pipettes. The resulting cell suspension was plated onto 22 mm glass coverslips previously coated with poly-L-ornithine (Sigma, mw 100,000 – 200,000), in a plating media consisting of: minimum essential medium (Gibco), 5 % v/v heat-inactivated horse serum (New Zealand origin, Gibco), 5 % v/v heat-inactivated fetal calf serum (South American origin, Gibco), 2 mM glutamine (Gibco), 20 mM glucose (Sigma), 100 u/100 µg /ml penicillin-G/streptomycin (Sigma). One hour after plating, the plating media was removed and replaced with maintenance media consisting of: Neurobasal-A (Gibco), 0.5x B27 supplement (Gibco), 0.5x Glutamax (Gibco), 35 mM D-glucose (sigma), 100 u/ml penicillin-G, 100 µg/ml streptomycin. The maintenance media was topped up with 0.5 ml fresh media twice a week for each 35 mm culture dish.

Neurons were transfected after seven days *in vitro* using Effectene (Qiagen). For this 0.4 µg of the indicated GABA_A_R subunit DNA (in pRK5) and 0.4 µg of eGFP DNA (pEGFP-C1) were mixed with: 100 µl EC buffer, 3.2 µl enhancer, 10 µl Effectene, 600 µl maintenance media. This solution was mixed by vortexing and added dropwise to neurons maintained in fresh media. Two hours after the addition of the transfection media, it was removed and replaced with conditioned media harvested from the neurons prior to the transfection.

### Analysis and whole-cell voltage clamp recordings of synaptic currents

Coverslips were removed from the incubator between 20 - 22 days *in vitro* and were placed into a bath continuously perfused with Krebs solution. Transfected cells were identified by their eGFP fluorescence. Only principal cells with pyramidal cell-like morphologies were used for recording. Patch electrodes with resistances of 2.5 - 3.5 MΩ were filled with a CsCl-based internal solution consisting of: 140 mM CsCl, 2 mM NaCl, 10 mM HEPES, 5 mM EGTA, 2 mM MgCl_2_, 0.5 mM CaCl_2_, 2 mM Na-ATP, 0.5 mM Na-GTP, 2 mM QX-314, were used for the whole-cell recording configuration. Recordings were made with an Axopatch 200B amplifier. Cells were subjected to a holding potential of −60 mV and series resistance compensation of > 85 % was applied. Recordings were made with sampling rates of 20 kHz using a 4-pole low pass Bessel filter set at 5 kHz. The access resistance was assessed every 5 min and used as a barometer of cell viability. If it deviated by more than 25 %, the data from the intervening period was discarded. For the recording of spontaneous inhibitory postsynaptic currents (sIPSCs), 2 mM kynurenic acid was included in the bath solution to block glutamate receptor activity. Bath temperature was controlled either by working at a controlled ambient temperature (21 °C, as measured with a thermometer and thermocouple immersed in the bath solution), or by warming the bath solution with an inline heater to 37 °C (confirmed by insertion of a thermocouple into the bath).

For experiments involving drug pre-treatments, final drug concentrations were added to the maintenance media (see above) of the cultured neurons, and the dishes were subsequently returned to the cell incubator for the duration of the treatment (20 min). The coverslips were then removed and placed into the recording chamber. Krebs solution was washed over the coverslip for 25 min prior to recording to ensure washout of the drug. For recordings performed 24 hrs after GABA treatment, the treatment media was removed, and the cells were washed 5x in HBSS, followed by one 20 min wash in fresh maintenance media before being returned to conditioned media harvested from the cells prior to the treatment.

Spontaneous IPSCs were identified in 5 min recording epochs using Clampfit (version 10.7, Molecular Devices) in conjunction with a template pattern search (generated from examples of sIPSC rise profiles and peak currents recorded from control cultured hippocampal neurons at the same temperature). All recorded events were visually inspected. For the calculation of mean amplitudes, events were discarded if they displayed inflections on the rising phase. Events used for kinetic analysis were also discarded if they displayed any secondary peaks during the decay phase. Conversely, for the calculation of rise-rates, inter-event intervals, and the production of cumulative probability distributions, all detected events were included. Decay curves were fitted to average sIPSC waveforms using a standard two component exponential function in ClampFit. Weighted time constants were determined from:

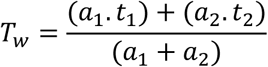

Where *a* indicates the relative weighting of each exponential component and *t* indicates the time constant of each component. Rise-time slopes were calculated using the maximum rate of rise function in ClampFit on data filtered at 2 kHZ (8-pole Bessel). For presentation purposes only, current records were further filtered with a 1 kHz 8-pole Bessel filter.

For peak-scaled non-stationary variance analysis, synaptic events were selected that displayed both clean rise and decay phases (without inflections, secondary peaks, or noise artefacts). These events were imported into WinWCP v5.2.3 (Dempster, 2015), and the peak of the averaged sIPSCs was aligned to the peaks of individual sIPSCs selected for analysis. The decay phases of individual sIPSCs were subtracted from the mean sIPSC decay to yield the current variance which was plotted against the corresponding mean current and the data curve fitted using the parabolic function:

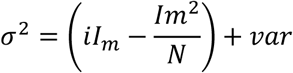

Where σ^2^ is the current variance, i is the single channel current, I_m_ is the mean current and N is the mean number of synaptic receptors activated during the peak of an sIPSC, and var is the baseline current variance. From the resulting parabolic curve fits, the equation was used to estimate both i and N for synaptic GABA_A_Rs.

### Cluster based analysis for sIPSCs

Spontaneous IPSC parameters were extracted using the Clampfit event detection protocol as described above. The following parameters were chosen for the clustering analysis: instantaneous sIPSC frequency, 10-90 % sIPSC rise time, 20-80 % rise time, 30-70 % rise time, maximum rise slope normalized to the peak sIPSC amplitude, rise slope normalized by the peak amplitude, inter-event sIPSC interval, burst window (calculated as the shortest interval containing 3 consecutive sIPSC events, including the one in question), and coefficient of variation within the burst window. Clustering of these parameters was carried out in R (ver 3.6.1), using the EMMIXskew package^72^ for analysing a mixture of multi-variate skew distributions. Skewed-t distributions^72^ were used to model the data, due to our observation that sIPSC parameters, such as frequency, often display a skew in their probability distributions. The number of clusters was established by running models with varying numbers of components, and then selecting the one which minimised the Bayes information criterion^73^. T-distributed stochastic neighbour embedding plots were produced in R using the Rtsne package^74^. All other plots were produced in OriginPro (2017). For the boxplots, boxes represent the interquartile range with the whiskers representing the 10-90 % interval.

### Immunoblotting

Dishes containing HEK cells were taken from the incubator and placed on ice. The media was removed, and the cells briefly washed in ice cold Tris-buffered saline (TBS). TBS was removed and radioimmunoprecipitation assay buffer (RIPA; 1 % NP-40, 0.5 % Na-deoxycholate, 0.1 % sodium-dodecyl sulfate (SDS), 150 mM NaCl, 50 mM Tris) was added. Dishes containing the cells and RIPA buffer were immediately placed into a −80 ^°^C freezer for approximately 5 min to freeze the cells. Plates were removed from the freezer, thawed on ice and cell lysates were collected in ice-cold Eppendorf tubes. Lysates were mixed by rotation at 4 °C for 1 hr to allow solubilisation of the membrane proteins to occur. The lysates were then spun at 13,000 rpm for 30 min. Bicinchoninic acid (BCA) assays (Thermo Scientific) were carried out and the protein concentrations normalised. Sample buffer was added to the lysate at final concentrations of: 17.5 % glycerol, 5 % SDS, 250 mM tris, 5 mg/ml bromophenol blue, 4 M urea, 200 mM dithiothreitol, and incubated at room temperature for 45 min prior to loading onto the gels.

Polyacrylamide gels (10 %) were prepared as follows: 10 % acrylamide, 375 mM bis-Tris, 0.2 % SDS, 0.15 % ammonium persulfate, 0.015% TEMED; and poured into glass cassettes (Biorad) and polymerised for at least 1 hr before sample loading. Samples were loaded directly onto the gel along with a pre-stained protein marker (New England Biolabs), and were run with a constant voltage of 60 V. Sodium metabisulfite was added to the electrophoresis running buffer (25 mM Tris, 192 mM glycine, 0.1% SDS) to a final concentration of 50 mM. Protein from the gel was then transferred to nitrocellulose membranes for 1 hr at 100 V. Blocking was next performed in TBS 0.1 % Tween (TBST) 5% BSA for 30 min at room temperature.

After blocking, proteins were exposed to primary antibodies overnight at 4oC (anti-γ2 phospho-S327, ab73183 (Abcam), 1:1000). Antibodies were diluted in the same buffer as used for blocking. Membranes were washed in TBST for 1 hr and probed with secondary antibodies (HRP-conjugate anti-rabbit (Rockland), 1:10000 in TBST 5 % BSA) for two hrs at room temperature. After exposure to the secondary antibody, the membranes were washed 3x in TBST (total wash time 1 hr) and were developed with luminata crescendo ECL substrate (Millipore) on an ImageQuant LAS 4000 (GE Healthcare). The membranes were stripped in mild stripping buffer (200 mM glycine, 0.1 % SDS, 1 % Tween20, pH 2.2), and subsequently re-probed for total γ2 GABA_A_R subunit levels (anti-γ2: Alomone, 1:500, AGA-005). Analysis was carried out using GelAnalyser 2010a. Lanes and bands were detected automatically, and a rolling-ball background correction was applied prior to quantification.

### Statistics

Unless otherwise indicated, all data are presented as means ± standard errors, with horizontal dashed lines representing the median value. Comparing means of multiple groups was undertaken using a one-way ANOVA followed by a Tukey post-hoc test. In the case of experiments with two conditions, two-tailed t-tests were performed. Results with p values < were defined as statistically significant (indicated by the * symbol on graphs), and p values are reported in full, where appropriate, within the text.

## Abbreviations

ANOVA: analysis of variance
Bis: bisindolylmalemide-I
CNS: central nervous system
Eto: etomidate
GABA: γ-aminobutyric acid
GABA_A_R: γ-aminobutyric acid receptor type A
HEK: human embryonic kidney cell line 293
IEI: inter-event interval
iLTP: inhibitory long-term potentiation
mIPSC: mini-inhibitory postsynaptic current
PKC: protein kinase C
PLC: phospholipase c
pLGIC: pentameric ligand-gated ion channel
PMA: phorbol 12-myristate 13-acetate
PS: pregnenolone sulfate
Ptx: picrotoxin
sIPSC: spontaneous inhibitory postsynaptic current
t-SNE: t-distributed stochastic neighbour embedding
TTX: tetrodotoxin

## Acknowledgements

This work was supported by an MRC programme grant (TGS). MF was supported by a 4-year MRC Postgraduate Studentship.

